# In-silico Targeting Phosphodiesterase 5 and In-vivo Evaluation of the Aphrodisiac Effect of *Azanza garckeana* Fruit

**DOI:** 10.1101/2024.10.30.621167

**Authors:** Ejiofor InnocentMary Ifedibaluchukwu, Ezeanya Jennifer Ifeoma, Ibe Chioma Ifeoma, Ogbue Onyeka Cyril, Emodi Rita Chisom, Agueze Jennifer Chinwendu, Ani Chibuike Sylvanus

**Affiliations:** Department of Pharmacognosy and Traditional Medicine, Faculty of Pharmaceutical Sciences Nnamdi Azikiwe, University, Awka, Anambra State, Nigeria; Department of Pharmaceutical and Medicinal Chemistry, Faculty of Pharmaceutical Sciences Nnamdi Azikiwe, University, Awka, Anambra State, Nigeria

**Keywords:** Aphrodisiac, sexual dysfunction, phosphodiesterase-5, molecular docking, *Azanza garckeana*

## Abstract

Sexual dysfunction is a persistent problem affecting men, with erectile dysfunction (ED) and premature ejaculation being the most common. The current treatment involving the use of PhPDE5 inhibitors, such as sildenafil, is faced with some serious side effects. In Nigeria, there are claims that *Azanza garckeana* fruit has positive effects in male sexual dysfunction. This study aims to evaluate this claim of the aphrodisiac effects of *Azanza garckeana*, using computational and animal models. The 3D structures of the protein target of interest (phosphodiesterase) and 21 phytocompounds already isolated from and identified in the fruit were obtained from the Protein Data Bank and PubChem, respectively. The protein targets and the phytocompounds were prepared for molecular docking simulation using the Autodock tool. The docking was performed using Autodock-vina on the Linux platform. Phytocompounds with high specificity for the protein targets were identified, and their solubility profile was obtained. The fruits were extracted with ethanol using a Soxhlet apparatus at 60°C and concentrated with a rotary evaporator at 40°C. Phytochemical tests were conducted on the extract, and acute toxicity was evaluated using OECD 423 guidelines. The aphrodisiac effect was studied in-vivo using a rat model. Parameters such as mount and intromission frequencies, mount and ejaculation latency and testosterone levels were assessed. Three phytocompounds showed better binding affinity for phosphodiesterase 5 than the reference compounds and were also found to be a good target for the proteins of interest. The extract was found to contain steroids, flavonoids, and terpenoids. The results from the in-vivo study demonstrated significant increases (*p*<0.05) in mean mounting and intromission frequencies and ejaculatory latency following treatment with the extract at 500 mg/kg body weight. The study also demonstrated a significant decrease (*p*<0.05) in mount latency. Testosterone levels increased significantly (*p*<0.05) at 500mg/kg body weight. This study affirms the aphrodisiac effect of *Azanza garckeana* fruit in the albino rat model and suggests that the fruit’s ethanol extract holds promise as an important phytomedicine for the development of more effective treatments for male sexual dysfunction.

## 1. Introduction

The intricate physiological process of male sexuality has a significant role in life quality. The human multi-system, which includes the neurological, cardiovascular, endocrine, and reproductive systems, must be coordinated to maintain proper sexual function [1–3]. The natural sex life will be compromised if the psychosocial aspects of the system above are altered. Male sexual dysfunction (SD) is a multifactorial illness. Sexual dysfunction refers to any difficulty encountered during any step of the male sexual process, including male sexual arousal, penile erection, penis insertion into the vagina, ejaculation, and any other obstacles [4,5]. The National Institutes of Health stated in 1992 that erectile dysfunction (ED) is the inability of the penis to attain or sustain the necessary firmness for fulfilling a fulfilling sexual life [6]. One of the most common and inadequately managed illnesses in SD is ED. Sexual function disorders are prevalent in men across all age groups and racial and cultural origins. According to the Massachusetts Male Aging Study, 52% of men aged 40 to 70 experience varied degrees of SD [7]. Despite a large body of basic research on SD, the pathogeny and risk factors of this illness are currently being investigated. Shaeer *et al* [8] investigated the prevalence of ED among males attending primary healthcare clinics in three countries with cultures substantially distinct from those of the industrialized West: Pakistan, Egypt, and Nigeria. This epidemiologic investigation, like that of Berrada et al [9], found that the prevalence of ED and other disorders linked with this condition is similar in Sub-Saharan Africa, the Middle East, and South Asia to that in the United States and Western Europe. Surveys of males aged 35 to 70 seeking primary medical care found that the age-adjusted prevalence of ED was 57.4% in Nigeria, 63.6% in Egypt, and 80.8% in Pakistan. As in other studies [10,11] older age, diabetes, prostate problems, and depression were all linked with increased risk for ED.

The current treatment involves the use of PDE5 inhibitors, such as sildenafil, tadalafil, and vardenafil, which are the first-line treatment for male erectile dysfunction.[12,13] PDE5 is an enzyme present in the smooth muscle of the corpora cavernosa that converts cGMP to 5’-GMP. As previously established, PDE5 inhibitors cause cGMP to accumulate, resulting in an inward flow of blood and a longer erection.[14,15,16,17] The most common side effects of PDE5 inhibitors are headaches, nasal congestion, flushing, dyspepsia, back discomfort, and myalgia.[13,15] Furthermore, PDE5 inhibitors can produce severe hypotension when taken with nitrates; consequently, patients taking nitrates should avoid PDE5 inhibitors. Steven-Johnson Syndrome, priapism, sudden vision loss, and sudden hearing loss are examples of rare adverse effects [13].

During sexual arousal, nitric oxide (NO) is released from nerve endings and endothelial cells in the corpus cavernosum. This NO stimulates guanylate cyclase, which converts guanosine triphosphate (GTP) into cyclic guanosine monophosphate (cGMP). The buildup of cGMP initiates a series of cGMP-dependent processes, resulting in the relaxation of smooth muscle in the corpus cavernosum and an increased flow of blood to the penis. PDE5 is an enzyme mainly located in the smooth muscle of the corpus cavernosum that specifically breaks down cGMP into 5′-GMP. PDE5 inhibitors, which have a structure similar to cGMP, bind competitively to the enzyme, preventing the breakdown of cGMP and amplifying the effects of nitric oxide (NO). The resulting increase in cGMP within smooth muscle cells helps maintain an erection for a longer duration [18].

The *Azanza garckeana* fruit (F. Hoffm.) Exell and Hillc, often referred to as Goron Tula (kola of Tula) in Hausa, are members of the Malvaceae family. It is solely grown in Gombe State’s Tula village in Nigeria. It is a versatile fruit from tropical Africa that can be eaten. In Northern Nigeria, it is a significant food and medicinal plant that is frequently used to make herbal remedies [19]. More than 20 different human diseases and ailments have supposedly been treated with *Azanza garckeana* in traditional medicine. Cough, chest pains, infertility, irregular menstruation, STDs, and liver impairments are among the illnesses for which the plant is utilized as a natural medicine [20,21]. *Azanza garckeana* has been shown to contain a variety of bioactive metabolites, such as amino acids, alkaloids, ascorbic acid, carotenoids, flavonoids, glucosides, phenols, lipids, tannins, and saponins [22].

In Nigeria, most especially in the Northern part of Nigeria, the fruit of this plant is used as an aphrodisiac. Previous studies on *Azanza garckeana* fruit revealed that the methanol and ethyl-acetate of fruit showed anti-oxidants, antimicrobial, anti-inflammatory, analgesic, anti-arthritic and wound healing effects [23]. The aqueous extract of *Azanza garckeana* fruit pulp has been shown to improve sexual behaviour and reproductive hormone concentrations, thereby potentially restoring sexual competence in sexually dysfunctional female rats. These findings provide additional support for the traditional use of *Azanza garckeana* in managing female sexual dysfunction [24]. In the present study, we utilized computation methods targeted at Phosphodiesterase 5 and an in-vivo approach to rationally evaluate the aphrodisiac effect of the *Azanza garckeana* fruit in male albino rats.

## 2. Materials and Methods

### 2.1 Identification of receptor/target

Literature was mined to determine the receptor/target involved in sexual dysfunction. This was conducted to assess the role of the receptors in the treatment of sexual dysfunction. This provides additional details regarding the receptor, its functions, and characteristics. Based on these pieces of information phosphodiesterase-5 was selected as a target for this study.

### 2.2 Identification of phytocompounds

Literature was reviewed to identify the Phytocompounds present in *Azanza garckeana* fruit.

### 2.3 Preparation of selected receptor/targets

The 3D structure of the target was obtained from the protein data bank (https://www.rcsb.org) with the pdb code 1XOZ (Phosphodiesterase 5). PyMOL tool was employed to gain insight into the protein and their co-crystallized ligands and edited. The protein was prepared for molecular docking simulation using AutoDockTools 1.5.6. Polar hydrogens were added and exported as a pdbqt (Protein Data Bank, Partial Charge (Q), & Atom Type (T)) file. Grid boxes were mapped around the active sites of the proteins individually and the XYZ coordinates were noted for sizes and centers at a grid space of 1.0 Å. The grid box parameters generated are shown in Table 1.

**Table 1.** Grid box parameters used in the molecular docking simulations.

|  | 1XOZ |  |
| --- | --- | --- |
|  | Center | Size |
| X | 47.364 | 26 |
| Y | 35.197 | 19 |
| Z | 12.951 | 25 |

### 2.4 Preparation of Ligands (Drug/Phytocompounds)

The phytocompounds in *Azanza garckeana* fruit and reference compounds were downloaded from PubChem in SDF-3D format. These reference compounds include compounds co-crystallized with the receptors on Protein Data Bank and other approved drugs effective against the selected targets. Utilizing AutoDockTools, the ligands were prepared for molecular docking simulations, including the assignment of all rotatable bonds, torsions, and force field minimization, and saved as pdbqt files. The reference compounds used include avanafil, sildenafil, tadalafil and vardenafil.

### 2.5 Validation of Docking Protocol

To validate the molecular docking simulations protocol for 1XOZ protein, the PDB structure of the protein in complex with its co-crystallized ligand was reproduced in-silico through molecular docking simulation which was executed using Autodock Vina [25] on a Linux platform based on the centers and sizes shown in table 1, with a virtual screening shell script. Docked conformation was then visualized in PyMol and poses were compared with the experimental crystal structure of the co-crystallized ligand.

### 2.6. Molecular Docking Simulation

The phytocompounds were batched for molecular docking simulations against 1XOZ. Molecular docking simulation was carried out in triplicates on a Linux platform using Autodock Vina after the validation of the docking protocol.

### 2.7 Post –Docking Analysis

The mean and ± SD (standard deviation) of the binding affinities after docking were calculated and recorded on a Microsoft Excel spreadsheet. The binding affinities of the phytocompounds were compared with the reference binding affinities. The phytocompounds with binding affinities lower than that of the reference drugs for each protein were selected.

### 2.8 Assessment of protein-ligand interaction

The interaction between the protein and ligand was assessed using the Discovery Studio visualizer. This assessment aimed to ascertain the exact amino acids that interacted with the ligands in comparison with the reference drugs.

### 2.9 Swiss Target Prediction

The Swiss target prediction was done to assess the targeting ability of the frontrunner phytocompounds in comparison to the reference drugs.

### 2.10. Solubility profiling

The solubility profile of the frontrunner phytocompounds was ascertained through literature mining.

### 2.11 Plant collection and identification

Fresh fruits of *Azanza garckeana* were collected from Abuja, Nigeria in November 2023. The fruit samples were identified and authenticated at the Department of Pharmacognosy and Traditional Medicine Faculty of Pharmaceutical Sciences, Agulu campus, Nnamdi Azikiwe University. The voucher number is NAU/PCG/AG024/011.

### 2.12 Preparation of Plant Sample

Fresh fruits of *Azanza garckeana* were purchased from Utako district market, Abuja. The fruits were washed under running tap water. The internal content of the fruits was scrapped out with the seeds using a spatula and transferred to a neat bowl.

### 2.13 Extraction of Crude Extract

The sample (855g) was extracted with ethanol using Soxhlet apparatus at 60℃ to completeness. The extract was concentrated in a rotary evaporator at 50℃ and then evaporated to dryness in a water bath at 50℃. The percentage yield was calculated using the formula stated below:

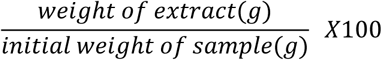

### 2.14 Phytochemical tests

The test was carried out according to the procedures outlined by Firdouse *et al*. 2011 [26] and Mahmud *et al*. 2015 [27].

### 2.15 Experimental Animals and Housing

Female mice and male albino rats weighing 18 to 22 grams and 150 to 200 grams were employed in the acute toxicity and in-vivo aphrodisiac evaluation studies respectively. The mice and rats were housed in animal houses with three animals per cage, maintained at 25 ± 2°C temperature, 50–60% humidity, and 12-hour light-dark cycles. Water and food were given to the mice. The Institution Animal Research Ethics Committee gave its approval before the start of the study (NAU/AREC/2024/0096).

### 2.16 Acute toxicity test

This acute toxicity study was conducted by OECD guideline 423 [28]. The test protocol entails giving a test sample at a high dose to a group animals of three (3) and monitoring them for signs of toxicity. If any symptoms of toxicity appear, the test is repeated using a lower dose of the test sample until a safe dose is achieved.

Female albino mice were used in the study. The animals were fed an oral initial dosage of 2000 mg of the extracts per kilogram of body weight, followed by a further two hours of fasting. Following dosage, each animal was observed alone for 30 minutes, then again every 24 hours, and finally every day for a total of 14 days. Changes in the animals’ weight, skin and fur, eyes, and mucous membranes, behaviour patterns, tremors, convulsions, salivation, diarrhoea, lethargy, sleep, coma, and death were all detected.

### 2.17 Experimental design for the in-vivo aphrodisiac effect

Male and female albino rats were used for this study. The *in-vivo* aphrodisiac potentials of *Azanza garckeana* fruit were studied in male rats using the method employed by Erhabor and Idu, 2017 [29]. The extract and the standard were administered as shown in table 2 for 14 days. On days 1, 7, and 14 of the extract administration, after 30 minutes, female rats were introduced into the cage in the ratio of 1:1 (male: female) and the male rat’s sexual behavioural parameters like mounting frequency, intromission frequency, mounting latency and ejaculatory latency were observed for 30 minutes. The mean values were calculated afterwards. Serum testosterone was monitored on day 14 of the treatment.

**Table 2.** Experimental design.

| Group | Number of animals | Administration | Group name |
| --- | --- | --- | --- |
| I | 5 | Water (10ml/kg) | Negative control |
| II | 5 | Sildenafil (50mg/kg) | Positive control |
| III | 5 | Crude extract (100mg/kg) | Extract administered in low dose (100mg/kg) |
| IV | 5 | Crude extract (250mg/kg) | Extract administered in medium dose (250mg/kg) |
| V | 5 | Crude extract (500mg/kg) | Extract administered in high dose (500mg/kg) |

### 2.18 Testosterone level analysis

The analysis of testosterone levels in the male rats used for the aphrodisiac study was analysed on day 14 of the study using a TETII kit. The TETII kit stored at 2-8℃ was removed from the refrigerator and used immediately. 100ul of blood sample was pipetted with a solid phase receptacle (SPR), the pipetting device, and the solid phase. The TETII SPR and TETII strips were inserted into the instrument. The assay was initiated following the instructions stated in the user’s manual. All the assay steps were performed automatically by the instrument. The assay was completed in 40 minutes and the SPR and strips were removed. The used SPR and strips were disposed of thereafter.

### 2.19 Statistical analysis

The data obtained from the study were analyzed using the Microsoft Excel package 2010. Results were expressed using mean, standard deviation, and percentage. One-way ANOVA and Tukey’s Honest Significant Difference Post Hoc Test were done on the VassarStats platform authored by Richard Lowry (http://vassarstats.net/).

## 3 Results

### 3.1 Identification of receptors/target

Presented in Figure 1 is the 3D representation of Phosphodiesterase 5 (1XOZ) as obtained from the protein data bank.

**Figure 1:**
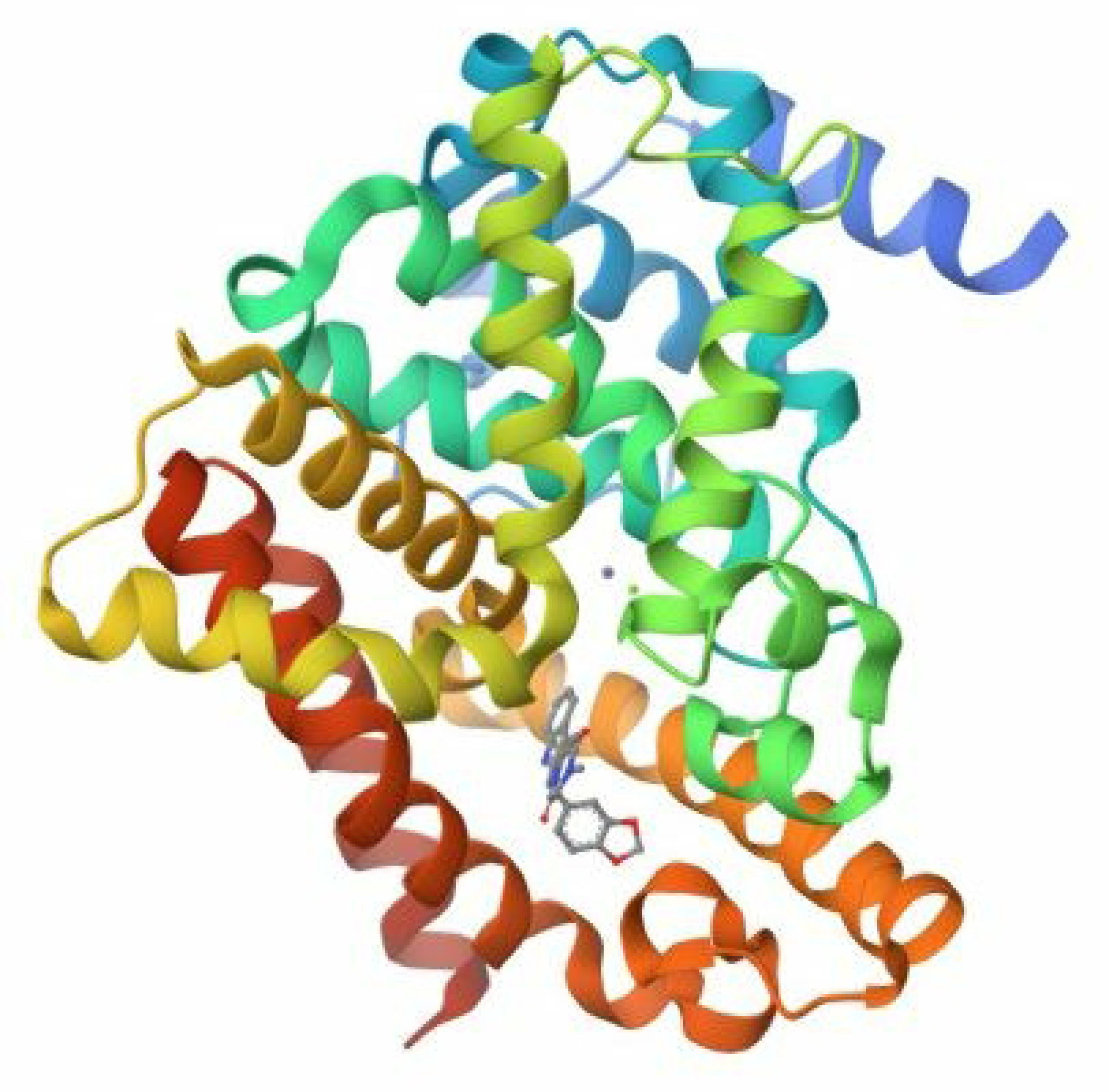
1XOZ as obtained from

### 3.2. Identification of phytocompounds

A total of 24 phytocompounds were found to have been identified in *Azanza garckeana* [30]. The phytocompounds include 6,6’-dimethoxygossypol, acrylate, arginine, aspartic acid, Azanzone, Azanzone B, Betulinic acid, Cadinane, Docosyl acrylate, Glutamic acid, Gossypol, Leucine, Lysine, Mansonone D, Mansonone E, Mansonone F, Mansonone G, Mansonone H, o-Naphthoquinone, Phenylalanine and Stigmasterol.

### 3.3. Validation of Docking Protocol

The result of the docking validation is presented in Figure 2. The docking validation was done for each of the targets’ co-crystallized ligands. In the figure, the green-coloured structure is the structure of the target co-crystallized ligand as obtained from the protein data bank, while the light blue-coloured structure is the docked ligand.

**Figure 2:**
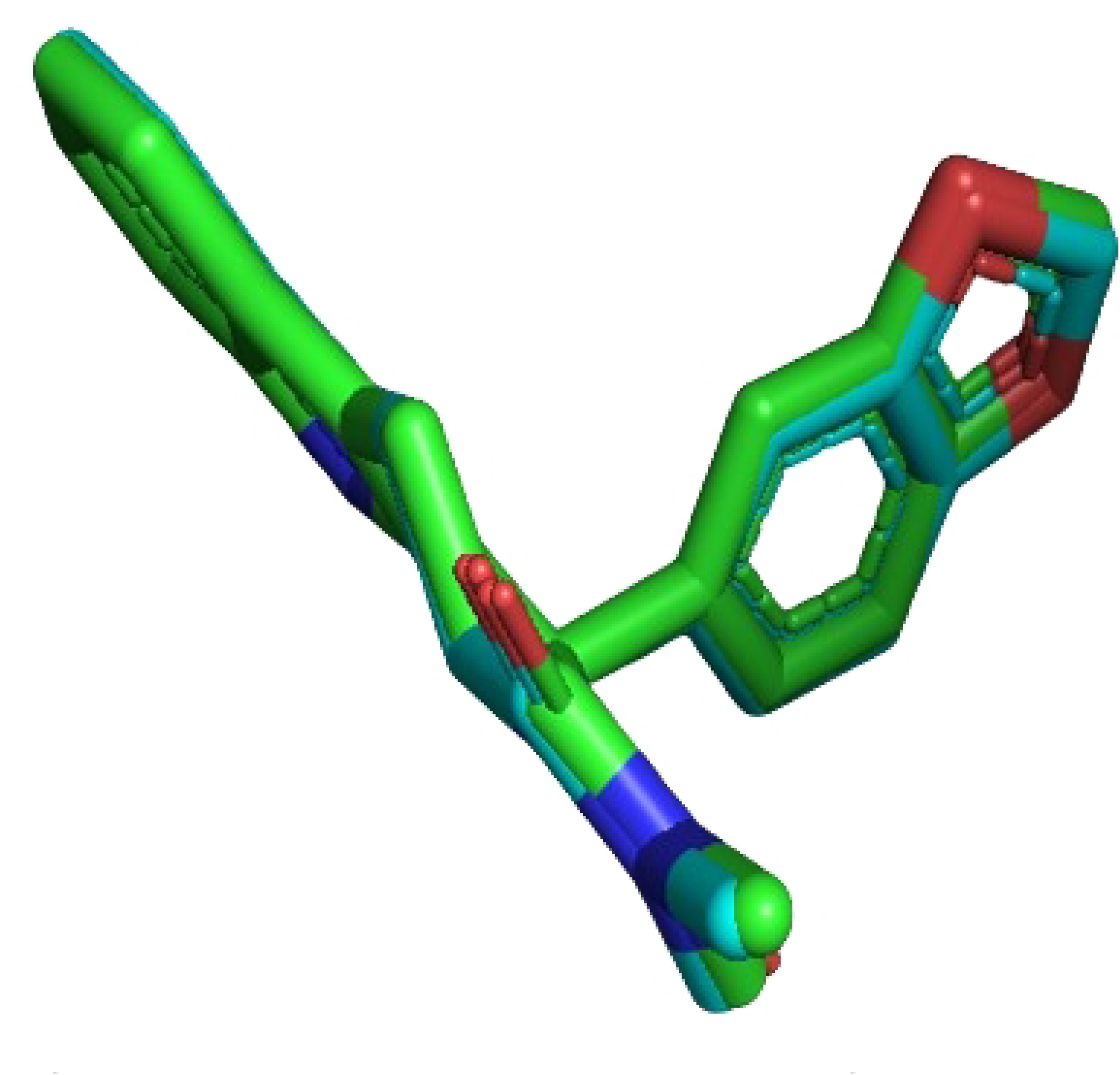
1XOZ superimpose

### 3.4. Molecular Docking Simulation

The result of the molecular docking simulation is presented in table 3. The phytocompounds presented in the table are the frontrunner phytocompounds. The frontrunner phytocompounds were determined based on the last reference drug with the highest value of binding affinity.

**Table 3.**
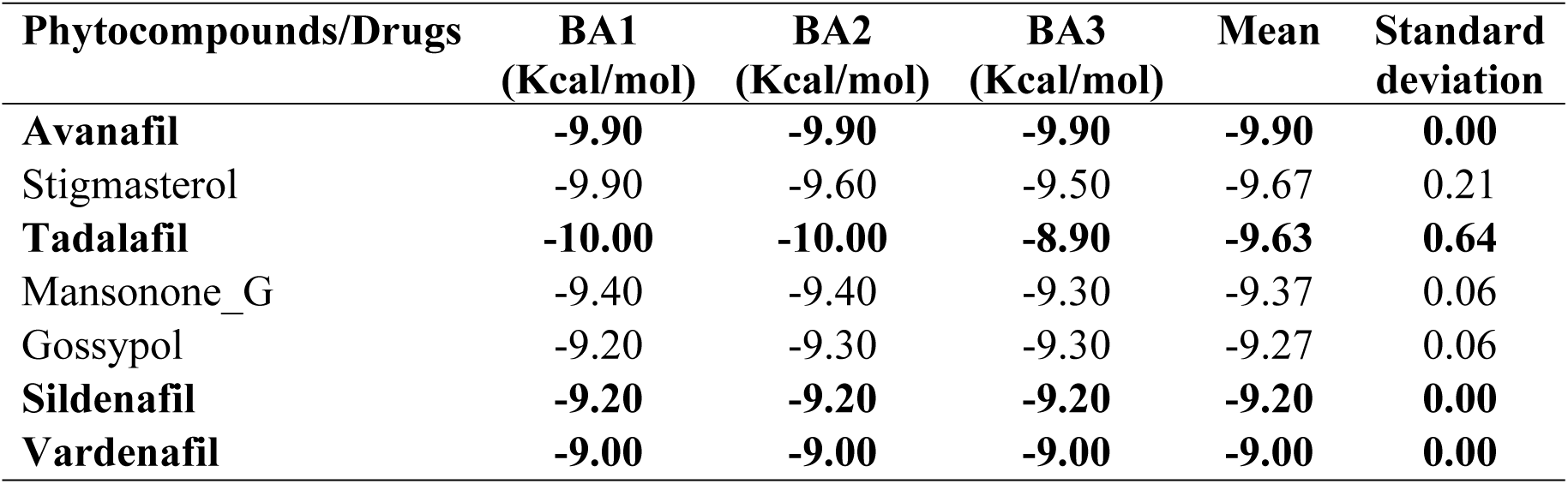
Frontrunner phytocompounds with low binding affinity (BA) and the reference drugs (in bold) for protein 1XOZ.

| Phytocompounds/Drugs | BA1<br>(Kcal/mol) | BA2<br>(Kcal/mol) | BA3<br>(Kcal/mol) | Mean | Standard<br>deviation |
| --- | --- | --- | --- | --- | --- |
| **Avanafil** | **-9.90** | **-9.90** | **-9.90** | **-9.90** | **0.00** |
| Stigmasterol | -9.90 | -9.60 | -9.50 | -9.67 | 0.21 |
| **Tadalafil** | **-10.00** | **-10.00** | **-8.90** | **-9.63** | **0.64** |
| Mansonone_G | -9.40 | -9.40 | -9.30 | -9.37 | 0.06 |
| Gossypol | -9.20 | -9.30 | -9.30 | -9.27 | 0.06 |
| **Sildenafil** | **-9.20** | **-9.20** | **-9.20** | **-9.20** | **0.00** |
| **Vardenafil** | **-9.00** | **-9.00** | **-9.00** | **-9.00** | **0.00** |

### 3.5. Assessment of Protein-Ligand Interaction

The results of the amino acid_drugs/phytocompounds interaction of the frontrunner phytocompounds and reference compounds with phosphodiesterase-5 are presented in Table 4. The assessment was done based on the 2D interaction observation of the interaction and taking notes of the amino acids that interacted with the frontrunner phytocompounds, in comparison to the co-crystalized ligand on the target protein as obtained from the protein data bank and also compared with the reference drugs.

**Table 4.** 1XOZ amino acid interaction with compounds.

| Drugs and<br>phytocompounds/Amino acids | ILE768 | ILE778 | ALA767 | VAL782 | PHE786 | LEU804 | MET816 | GLN817 |
| --- | --- | --- | --- | --- | --- | --- | --- | --- |
| Tadalafil (Co-crystallized ligand) | + | + | + | + | + | + | + | + |
| Avanafil | + | - | + | + | - | + | + | + |
| Sildenafil | - | - | + | + | + | - | - | + |
| Vardenafil | - | - | - | - | + | + | - | - |
| Gossypol | - | - | - | - | + | + | + | + |
| Mansonone_G | + | - | - | + | - | + | - | + |
| Stigmasterol | + | - | + | + | + | - | - | - |

### 3.6. Solubility profiling

The solubility profile of the frontrunner compounds is presented in Table 5. From the result, it can be observed that ethanol and chloroform were regular for the solubility of the phytocompounds. For this present study, we selected ethanol for extraction.

**Table 5.** Solubility profile of the frontrunner phytocompounds.

| Phytocompounds | Solubility |
| --- | --- |
| Cadinane | Ethanol, ether, chloroform, hexane |
| Gossypol | Ethanol, methanol, chloroform, ether |
| Mansonone D | Ethanol, methanol, chloroform ether |
| Mansonone E | Chloroform, ether, benzene, ethanol |
| Mansonone G | Ethanol, methanol, chloroform, ether |
| Mansonone H | Ethanol, methanol, chloroform ether |
| Stigmasterol | Chloroform, ether, benzene, ethanol |
| o-naphthoquinone | Methanol, acetone, chloroform, ethanol |
| 1,4-naphthalenedione | Acetone, methanol, ethanol, chloroform |

### 3.7. Extractive yield

The extractive yield was calculated using the following parameters;

Weight of the sample = 855g

Weight of the dried extract = 110

Percentage yield = 12.8 %

### 3.8. Phytochemical tests

Presented in Table 6 is the result of the qualitative phytochemical tests, showing the class of the phytochemicals present in the extract. The positive sign represents the phytochemicals present, while the negative sign represents the phytochemicals absent.

**Table 6.**
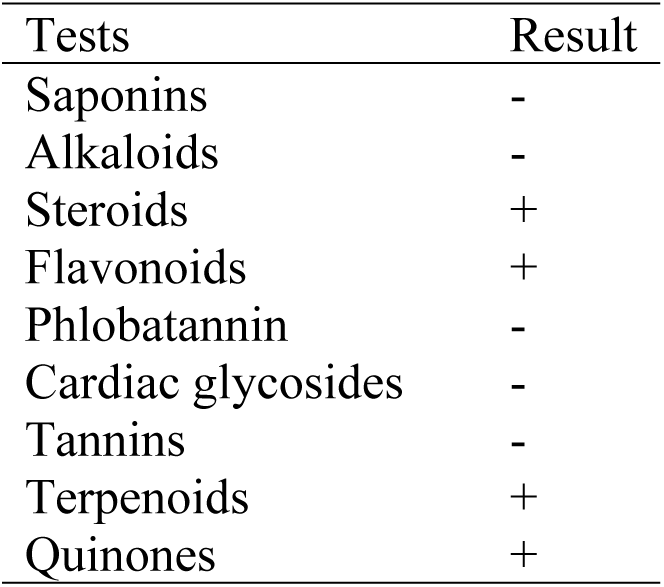
Phytochemical tests.

| Tests | Result |
| --- | --- |
| Saponins | - |
| Alkaloids | - |
| Steroids | + |
| Flavonoids | + |
| Phlobatannin | - |
| Cardiac glycosides | - |
| Tannins | - |
| Terpenoids | + |
| Quinones | + |

### 3.9. Acute toxicity test

From our observations during and after the acute toxicity test, none of the mice showed any observable signs of acute toxicity at 2000 mg/kg body weight.

### 3.10. Aphrodisiac effect analysis

#### 3.10.1. Mount frequency

Table 7 and Figure 3 show the mounting frequency, or the number of mounts that occur before ejaculation. The mounts were measured on days 1, 7, and 14 of the study. The number of mounts recorded increased in a dose-dependent manner throughout the research.

**Figure 3:**
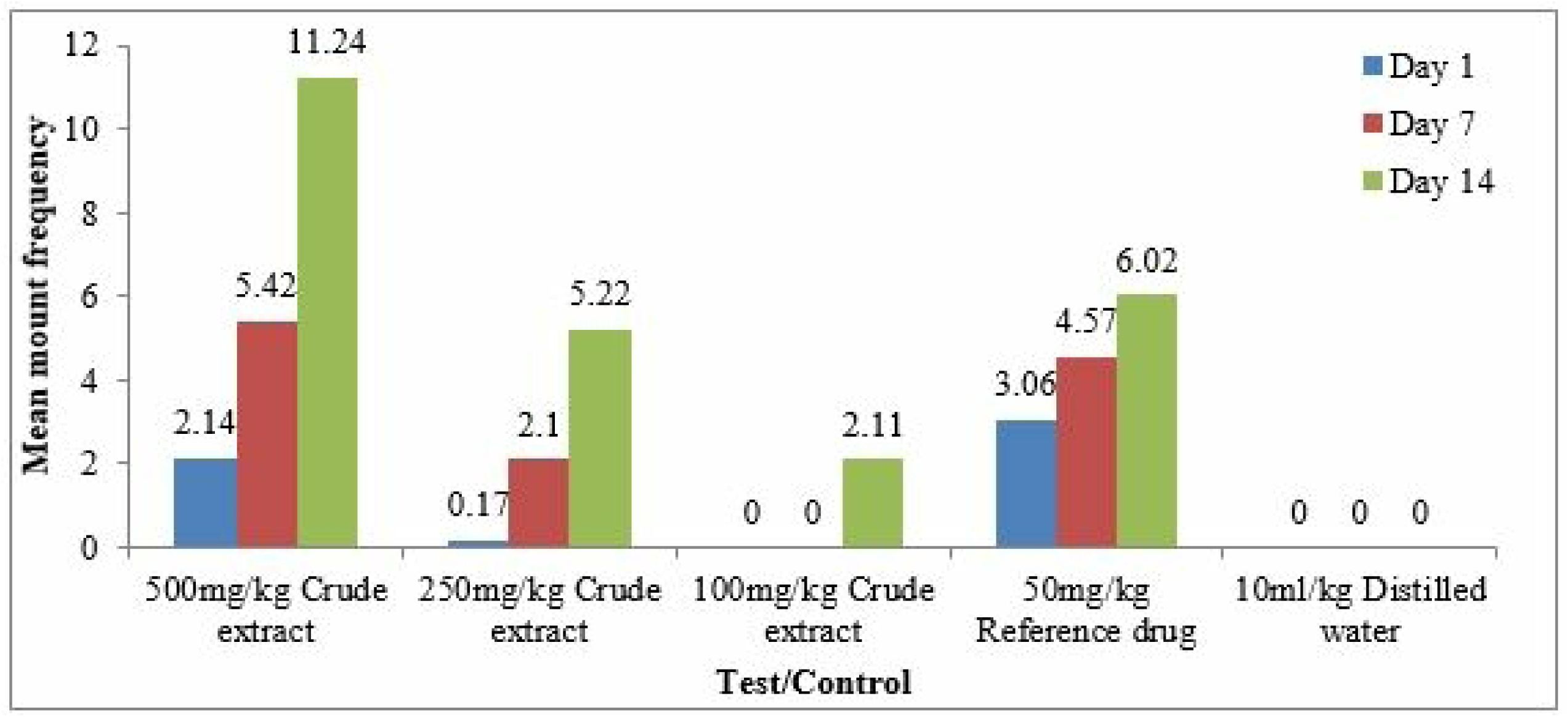
Chart of mean mount frequency

**Table 7.** Mean mount frequency.

| Treatment | Day 1 | Day 7 | Day 14 |
| --- | --- | --- | --- |
| 500mg/kg Crude extract | 2.14±0.08* | 5.42±0.20* | 11.24±0.30* |
| 250mg/kg Crude extract | 0.17±0.02* | 2.10±0.03* | 5.22±0.22* |
| 100mg/kg Crude extract | 0.00±0.00 | 0.00±0.00 | 2.11±0.06* |
| 50mg/kg Reference drug | 3.06±0.09 | 4.57±0.05 | 6.02±0.21 |
| 10ml/kg Distilled water | 0.00±0.00 | 0.00±0.00 | 0.00±0.00 |

Values are means of n = 5 (±SEM). Statistical significance for the extracts treated \* $p < 0.05$ compared to the control

#### 3.10.2. Mount latency

Table 8 and Figure 4 illustrate the mounting latency, which is the time between the addition of a receptive female into the arena and the first mount. Days 1, 7, and 14 of the investigation were used to record the mounts. Throughout the research, the latency decreased in a dose-dependent manner.

**Figure 4:**
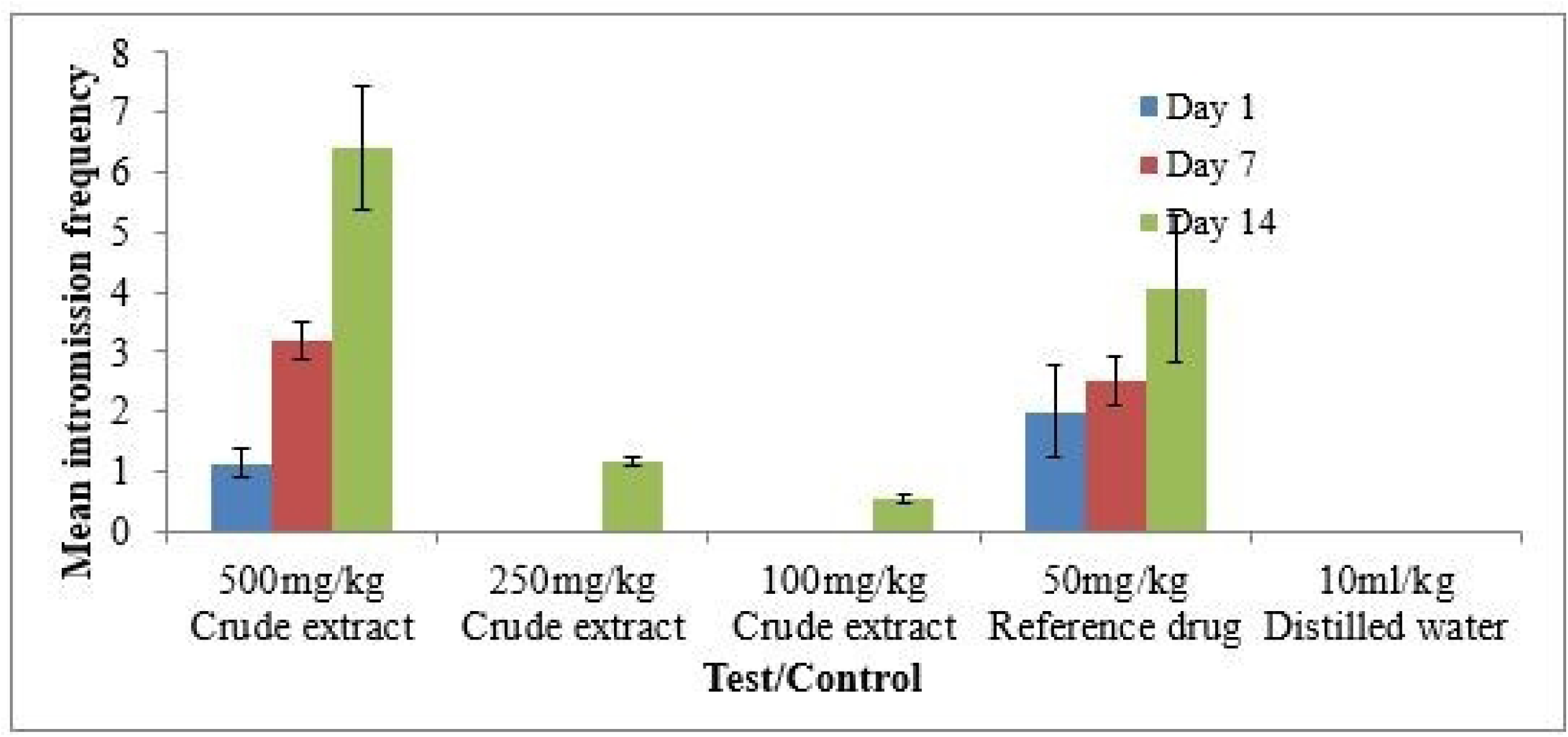
Chart of mean intromission frequency

**Table 8.** Mean mount latency (in minutes)

| Treatment | Day 1 | Day 7 | Day 14 |
| --- | --- | --- | --- |
| 500mg/kg Crude extract | 2.08±0.61* | 1.03±0.32* | 0.17±0.07* |
| 250mg/kg Crude extract | 0.00±0.00 | 0.00±0.00 | 3.06±0.71 |
| 100mg/kg Crude extract | 0.00±0.00 | 0.00±0.00 | 7.23±1.02 |
| 50mg/kg Reference drug | 1.32±0.05 | 1.04±0.12 | 0.96±0.06 |
| 10ml/kg Distilled water | 0.00±0.00 | 0.00±0.00 | 0.00±0.00 |

Values are means of n = 5 (±SEM). Statistical significance for the extracts treated \* $p < 0.05$ compared to the control

#### 3.10.3. Intromission frequency

Table 9 and Figure 5 illustrate the intromission frequency, which is the total number of intromissions from when the female animal was introduced until ejaculation. Days 1, 7, and 14 of the investigation were used to record the mounts. Throughout the research, intromissions increased in a dose-dependent manner.

**Figure 5:**
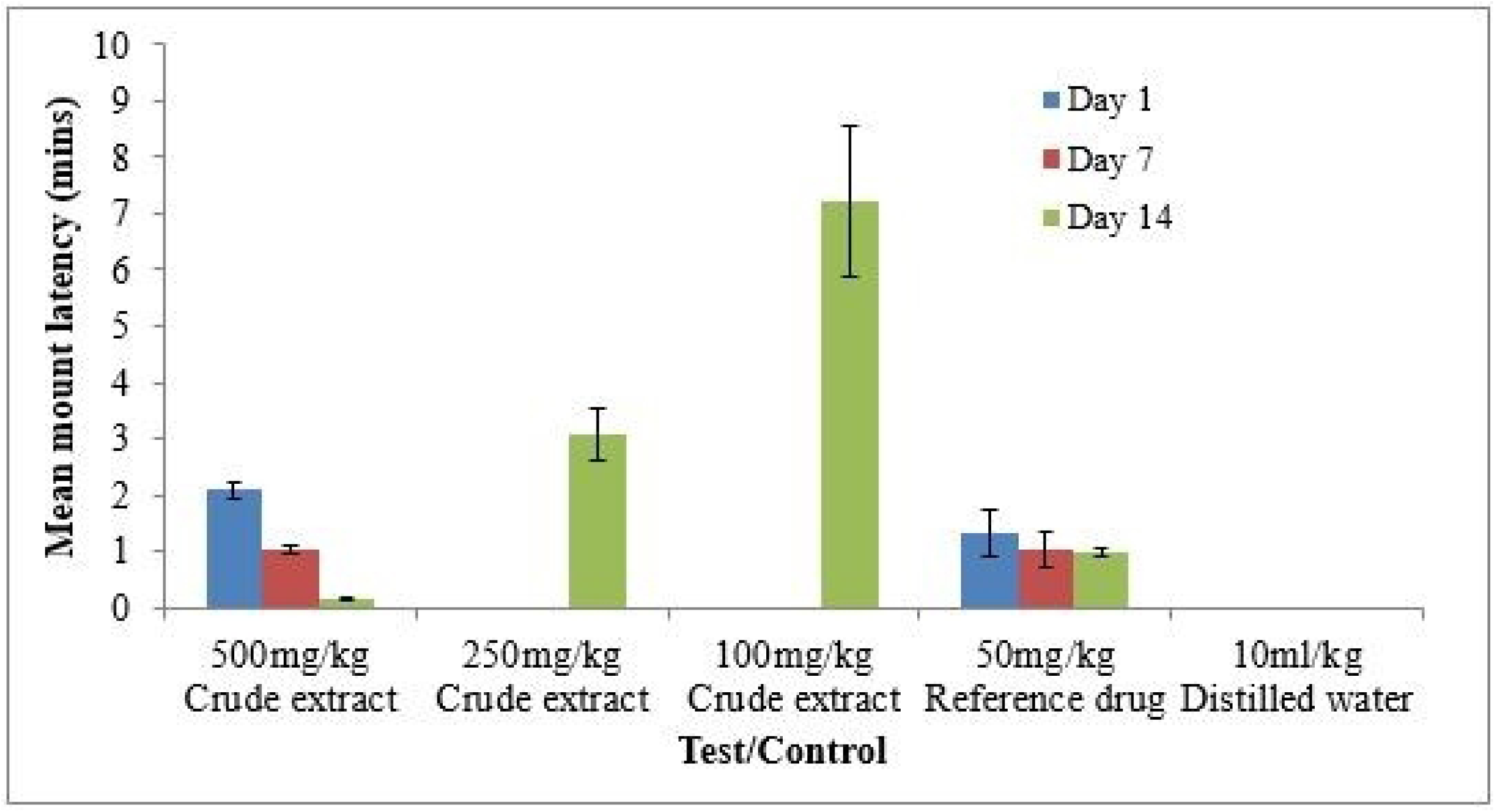
Chart of mean mount latency

**Table 9.** Mean intromission frequency.

| Treatment | Day 1 | Day 7 | Day 14 |
| --- | --- | --- | --- |
| 500mg/kg Crude extract | 1.14±0.30* | 3.21±1.06* | 6.40±1.60* |
| 250mg/kg Crude extract | 0.00±0.00 | 0.00±0.00 | 1.17±0.56* |
| 100mg/kg Crude extract | 0.00±0.00 | 0.00±0.00 | 0.53±0.05* |
| 50mg/kg Reference drug | 2.01±0.83 | 2.53±0.63 | 4.07±0.15 |
| 10ml/kg Distilled water | 0.00±0.00 | 0.00±0.00 | 0.00±0.00 |

Values are means of n = 5 (±SEM). Statistical significance for the extracts treated \* $p < 0.05$ compared to the control

#### 3.10.4. Mean ejaculatory latency

The ejaculatory latency, or the amount of time (measured in minutes) between the first intromission and ejaculation, is shown in Table 10 and Figure 6. The study employed days 1, 7, and 14 to document the ejaculatory delay. Ejaculatory delay rose in a dose-dependent way throughout the study.

**Figure 6:**
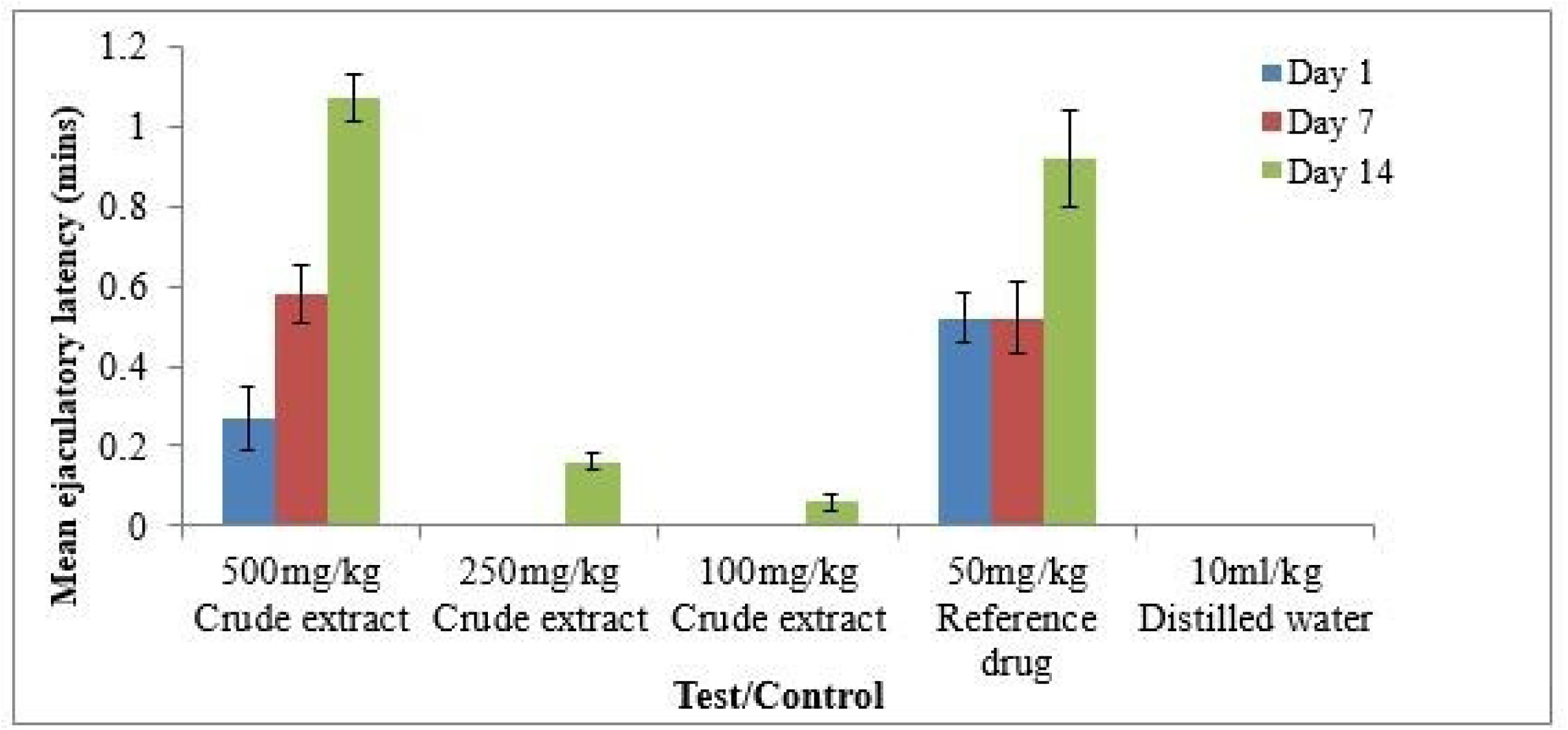
Chart of mean ejaculatory latency

**Table 10.** Mean ejaculatory latency.

| Treatment | Day 1 | Day 7 | Day 14 |
| --- | --- | --- | --- |
| 500mg/kg Crude extract | 0.27±0.08 | 0.58±0.05* | 1.07±0.10* |
| 250mg/kg Crude extract | 0.00±0.00 | 0.00±0.00 | 0.16±0.01 |
| 100mg/kg Crude extract | 0.00±0.00 | 0.00±0.00 | 0.06±0.02 |
| 50mg/kg Reference drug | 0.52±0.04 | 0.52±0.11 | 0.92±0.21 |
| 10ml/kg Distilled water | 0.00±0.00 | 0.00±0.00 | 0.00±0.00 |

Values are means of n = 5 (±SEM). Statistical significance for the extracts treated \* $p < 0.05$ compared to the control

### 3.11. Chemical parameters

The results of the testosterone levels of the test animals are presented in Table 11 and Figure 7. There was a dose-dependent increase in the levels of the testosterone.

**Figure 7:**
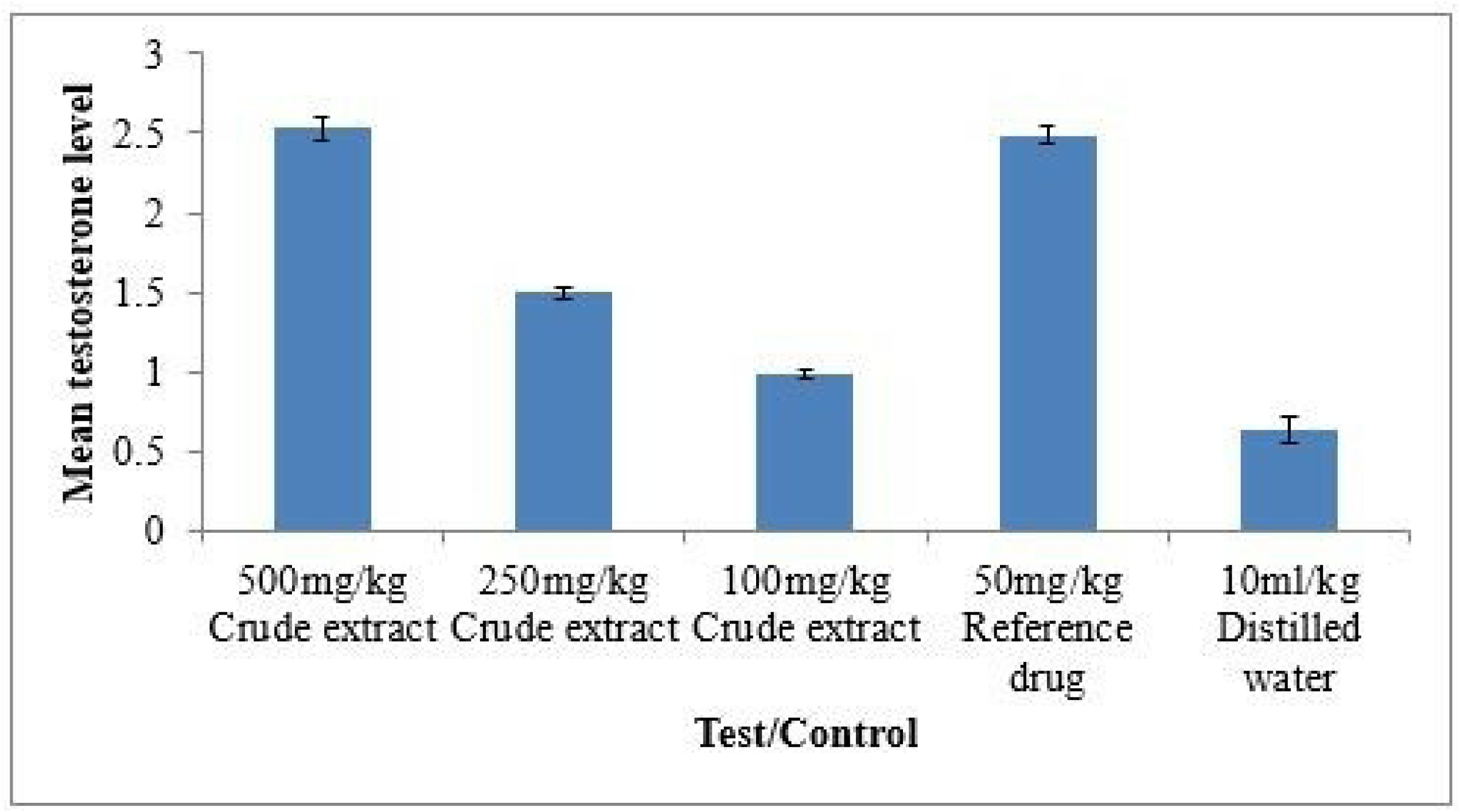
Chart of mean testosterone

**Table 11.** Testosterone levels.

| Treatment | Testosterone (ng/ml) |
| --- | --- |
| 500mg/kg Crude extract | 2.53±0.18* |
| 250mg/kg Crude extract | 1.50±0.09* |
| 100mg/kg Crude extract | 0.98±0.11* |
| 50mg/kg Reference drug | 2.49±0.05 |
| 10ml/kg Distilled water | 0.64±0.07 |

Values are means of n = 5 (±SEM). Statistical significance for the extracts treated \* $p < 0.05$ compared to the control

## 4. Discussion

Traditional medicine has long been known to be practised by Africans. The fruit *Azanza garckeana* has been used traditionally in Africa and beyond to treat sexual dysfunction. The most prevalent problems associated with sexual dysfunction include decreased sexual desire, erectile dysfunction, and disorders of ejaculation and orgasm [31]. This study evaluated and validated the claim of the high efficacy of *Azanza garckeana* in the treatment of sexual disorders. This study started with the in-silico evaluation of the aphrodisiac effects of *A. garckeana* phytocompounds, targeting both male and female sexual dysfunction, which necessitated our selection of serotonin and phosphodiesterase targets. The in-silico studies started with the validation of the molecular docking parameters. Several papers have been published emphasizing the need for initial validation of the docking protocols and sufficient hardware information for molecular docking studies [32,33]. Re-docking is a common method used to assess the precision of docking processes. Re-docking is a widely used method for determining a docking program’s accuracy for the first time. It replicates the co-crystallized ligand in its first posture as a validation technique. As was observed in Figure 2, the docking technique and parameters used were able to replicate the co-crystalized ligands as obtained from the protein data bank.

The molecular docking results presented in Table 3 were ranked based on their binding affinities expressed in Kcal/mol. In molecular docking simulation, the kind of contact between ligand and protein is indicated by the mode of interaction that has the lowest or most negative binding affinity. Also in molecular docking simulation, energy is released as a result of the interaction between the ligand and protein. As a result, the binding energy is shown as a negative value [34]. Stigmasterol, Mansonone_G, and Gossypol were identified as frontrunner phytocompounds because they have lower binding energies than two of the reference compounds, Sildenafil and Vardenafil.

The references and frontrunner phytocompounds interaction with the amino acids present in the protein active site shown in Table 4 was assessed using the Discovery studio visualizer. Along with the type of contact and bond lengths, the Discovery Studio aids in the identification of interactions between the active sites in the target and ligand/drug conformation. Discovery Studio is one of the distinctive, centralized, and user-friendly graphical interface for effective protein modelling and drug design [35,36]. For there to be a pharmacological activity, the ligands or drugs have to interact with specific amino acids in the target sites of a protein. In Table 4, the amino acids’ interaction with reference drugs was compared with those of the frontrunner phytocompounds. Observation revealed that amino acid interactions found in the reference drugs are also present in the frontrunner phytocompounds at different ranges.

The phytochemical test conducted after the extraction revealed some important information about the nature of the phytocompounds present in the extract. The phytocompounds targeted, are shown in the solubility profile of the frontrunner. Gossypol is a polyphenol with a yellow pigment similar to flavonoids [37]. Mansonones and o-naphthoquinone are quinones [38]. Stigmasterol is a steroid [39]. Most of the phytochemicals in Table 5 belong to the class of the phytochemicals present in the extract as shown in Table 6, which means that the solvent selection for extraction fulfilled its rational and target purpose. Even though further analytical studies are required to establish the presence of the specific frontrunner compounds in the extract.

The increase in mount frequency, intromission frequency, ejaculatory latency, and decrease in mount latency observed in the study in comparison with the control groups give insight into the aphrodisiac effect of the extract. While the number of mounts indicates sexual drive, a rise in intromissions indicates the effectiveness of an erection, penile orientation, and the ease with which ejaculatory reflexes are engaged. Both mount and intromission frequencies are useful indicators of vigour, libido, and potency [40,41]. Aphrodisiac properties are suggested by the mere prolongation of the ejaculatory latency. No suppression of mount-and-intromission frequency, copulatory effectiveness, or intercopulatory interval was seen in any of the treated rats that mounted and were introduced. This implies that the aphrodisiac activity did not impact desire, sexual vigour, or sexual performance. Sexual motivation is indicated by mount latency [42]. As a result, the study’s findings regarding the decrease in mount latency may indicate an increase in arousal and sexual drive. Further evidence for the extract’s ability to boost sex may come from the male rats’ increased sexual appetitive behaviour. The extracts and regular medication may have extended the duration of coitus, which is a symptom of increased sexual drive, based on the considerable increase in ejaculation latency [43]. The animals’ improved copulatory performance is indicated by the extract of *Azanza garckeana*’s longer ejaculation latency. The extracts of *Moringa oleifera* [44], *Allium tuberosum* [45], and *Monsonia angustifolia* [46] have all shown results that are comparable.

The increase in testosterone levels following the administration of the *Azanza garckeana* extract as observed in Table 11 and Figure 7, can also be associated with the aphrodisiac effect of the extract. An increase in anterior pituitary hormone and blood testosterone levels may have contributed to the enhancement of libido. The production of dopamine, a crucial neurotransmitter in the regulation of locomotor activity necessary for the presentation of copulatory and sexual behaviour, is thought to be stimulated by these hormones [40,47,48].

The considerable and long-lasting increase in sexual activity and aphrodisiac properties inherent in the extract is indicated by the notable increase in calculated parameters seen in the rats given the extract. Vasodilation, testosterone elevation, gonadotropin production, and nitric oxide production are some of the mechanisms thought to underlie the efficacy of plants used as aphrodisiacs [40,49]. In comparison to sildenafil, which was utilized as the reference in this investigation, treatment of the male rats with the *Azanza garckeana* extract improved their sexual behaviour.

## 5. Conclusion

The male albino rats used in this study showed dose-dependent sexual stimulatory effects from the *Azanza garckeana* fruit ethanolic extract, especially at the medium and the high dose used for the study. The findings support the indigenous people of northern Nigeria’s use of the *Azanza garckeana* fruit to manage male erectile dysfunction, premature ejaculation, and desire/libido issues. This study will also be replicated in female rats to observe its aphrodisiac effect in female animals. Further studies will be carried out to establish the phytocompounds present in the ethanol extract. Also chloroform extract also be evaluated for phytocompounds and aphrodisiac effects in the male and female rats.

## Acknowledgements

The authors wish to acknowledge Nnamdi Azikiwe University Awka and School of Pharmacy Agulu for the use of their facilities and equipment to conduct the experiments. The authors also appreciate the support of the CURIES research team, Faculty of Pharmaceutical Sciences, Nnamdi Azikiwe University, Awka, Anambra State, Nigeria.

